# Evaluation of a pan-*Leishmania* SL-RNA qPCR assay for parasite detection in laboratory-reared and field-collected sand flies and reservoir hosts

**DOI:** 10.1101/2020.01.08.898411

**Authors:** Myrthe Pareyn, Rik Hendrickx, Nigatu Girma, Sarah Hendrickx, Lieselotte Van Bockstal, Natalie Van Houtte, Simon Shibru, Louis Maes, Herwig Leirs, Guy Caljon

**Affiliations:** Biology department, University of Antwerp, Wilrijk, Belgium; Laboratory of Microbiology, Parasitology and Hygiene, University of Antwerp, Wilrijk, Belgium; Biology department, Arba Minch University, Arba Minch, Ethiopia

## Abstract

**Background:** In eco-epidemiological studies, *Leishmania* detection in vectors and reservoirs is frequently accomplished by high-throughput and sensitive molecular methods that target minicircle kinetoplast DNA (kDNA). A pan-*Leishmania* SYBR Green quantitative PCR (qPCR) assay which specifically detects the conserved spliced-leader RNA (SL-RNA) sequence has recently been developed. This study comparatively assessed the SL-RNA assay performance for the detection of *Leishmania* in field and laboratory infected sand flies and in tissue samples from hyraxes as reservoir hosts.

**Principal findings:** The qPCRs targeting SL-RNA and kDNA performed equally well on infected sand fly samples, despite preservation and extraction under presumed unfavorable conditions for downstream RNA detection. Nucleic acid extraction by a crude extraction buffer combined with a precipitation step was highly compatible with downstream SL-RNA and kDNA detection. Copy numbers of kDNA were found to be identical in culture-derived parasites and promastigotes isolated from sand fly midguts. SL-RNA levels were approximately 3-fold lower in sand fly promastigotes (ΔCt 1.7). The theoretical limit of detection and quantification of the SL-RNA qPCR respectively reached down to 10^−3^ and 10 parasite equivalents. SL-RNA detection in stored hyrax samples was less efficient with some false negative assay results, most likely due to the long-term tissue storage in absence of RNA stabilizing reagents.

**Conclusion:** This study shows that a crude extraction method in combination with the SL-RNA qPCR assay is suitable for the detection and quantification of *Leishmania* in sand flies. The assay provides complementary information to the standard kDNA assays, since it is pan-*Leishmania* specific and detects viable parasites, a prerequisite for identification of vectors and reservoirs.

**Author summary:** In order to identify vectors and reservoirs of *Leishmania*, a large number of sand fly and animal tissue samples needs to be screened, because the infection prevalence is generally low. Hence, sensitive low-cost methods are required for nucleic acid isolation and *Leishmania* detection. Most approaches amplify DNA targets, in particular minicircle kinetoplast DNA (kDNA). Recently, a qPCR was developed that detects the spliced-leader RNA (SL-RNA) sequence, which is conserved among various *Leishmania* species and allows detection of viable parasites. We show that the SL-RNA qPCR is highly compatible with a low-cost, crude extraction approach and performs equally well on laboratory and field infected sand fly samples as kDNA qPCR assays. The assay can detect 10^−3^ parasite equivalent in sand flies and enables *Leishmania* quantification down to 10 parasites. We found that the copy number of SL-RNA is 3-fold lower in sand fly derived promastigotes compared to cultured promastigotes. SL-RNA detection in hyrax tissue samples appeared less efficient, which is presumably due to long-term storage without RNA stabilizing reagents. Overall, our assay is complementary to kDNA assays as it can identify viable *Leishmania* stages, which provides pivotal information for identification of reservoirs and vectors and their transmission capacity.

## Introduction

Leishmaniasis is a vector-borne disease caused by protozoa of the genus *Leishmania*, which are transmitted during the blood feeding of female phlebotomine sand flies. The infection can be manifested in three major clinical forms: cutaneous (CL), mucocutaneous (MCL) and visceral (VL) leishmaniasis [1]. In Ethiopia, *L. aethiopica* is the predominant species causing CL and its vectors are *Phlebotomus longipes* and *P. pedifer* [2–4]. Hyraxes (*Heterohyrax brucei* and *Procavia capensis*) have been found asymptomatically infected with *L. aethiopica* in large numbers, indicating that they are major animal reservoirs in Ethiopia [3–5].

For eco-epidemiological research, there is a need for sensitive, high-throughput methods to identify and quantify *Leishmania* parasites in (potential) vectors and hosts [6]. The golden standard for parasite detection in sand flies and animal tissues is microscopic examination. This method allows to confirm the presence of viable parasites, but is time consuming and requires a substantial level of expertise [7]. These drawbacks resulted in a shift towards sample screening with molecular assays. Procedures generally start with nucleic acid extraction for which efficient, but expensive kits are commercially available. Low-cost methods, like organic (*i.e.* phenol-chloroform) or chelex extractions, are widely utilized, but have disadvantages. The former method is very time consuming and often involves toxic chemicals while the latter only yields low amounts of genomic DNA [8]. Extraction approaches with lysis buffers containing SDS, EDTA, Tris-HCl and NaCl have been applied to various tissues [9], although this crude procedure may lead to inhibition in downstream molecular applications [8].

A variety of (real-time) PCR methods targeting different gene fragments has been described, many of which remain to be validated on multiple *Leishmania* species and different tissues, or have issues regarding quantification [10,11]. The most commonly used PCR assay for *Leishmania* detection in sandflies [12,13] and small mammals [14–16] is targeting the minicircle kinetoplast DNA (kDNA). Because of the high kDNA copy number (10^4^ minicircles per parasite), very low numbers of parasites can be detected [7]. However, the nucleotide sequence and copy number sometimes differ among *Leishmania* species, impeding consistent quantification [17,18]. Another concern is that it sometimes results in false positive assay results due to its high sensitivity, even though all preventive measures to avoid contamination are taken [19–21].

Few studies investigated the use of RNA targets for parasite detection, although these may be more informative than DNA targets given the ability to discriminate viable parasites [22]. Recently, a pan-*Leishmania* SYBR Green quantitative PCR (qPCR) assay has been developed, targeting the highly conserved mini-exon encoded 39 bp spliced-leader RNA (SL-RNA) sequence, which shows excellent sensitivity and specificity. The assay was able to detect eight Old- and New-World *Leishmania* species with equal threshold cycle (Ct) values and was validated on tissue samples of *L. infantum* infected hamsters, promastigote spiked human blood and blood nucleic acid extracts from visceral leishmaniasis patients. It appeared that the limit of detection (LoD) of the SL-RNA qPCR was one log lower than the LoD of a Taqman multiplex assay targeting kDNA [23].

In this study, we aimed to evaluate the SL-RNA qPCR assay in combination with a crude extraction procedure for detection and quantification of *Leishmania* parasites in field- and laboratory-collected (infected) sand flies and hyrax tissue samples collected in Ethiopia.

## Materials and methods

### Ethics statement

The used chicken skins were obtained from day-old male chicks of a layer breed (Verpymo, Poppel, Belgium). The euthanasia of the chicks and use of laboratory rodents were carried out in strict accordance with all mandatory guidelines (EU directives, including the Revised Directive 2010/63/EU on the Protection of Animals used for Scientific Purposes that came into force on 01/01/2013, and the declaration of Helsinki in its latest version). All animal handlings were approved by the Ethics Committee of the University of Antwerp, Belgium (UA-ECD 2016–54 (2-9-2016)).

Hyrax trapping and sample collection in Ethiopia were conducted with authorization of the appropriate institutional authorities. Handling of the animals was carried out according to the 2016 Guidelines of the American Society of Mammalogists for use of small mammals in research and education.

### Parasites

The *L. major* strain MHOM/SA/85/JISH118 used in this study was cultivated *in vitro* at 26°C in HOMEM promastigote medium (Gibco, Life Technologies, Belgium), supplemented with 10% inactivated fetal calf serum (Invitrogen, Belgium) and was sub-cultured twice weekly.

### Sand flies

*Lutzomyia longipalpis* sand flies were maintained at the insectary of the Laboratory of Microbiology, Parasitology and Hygiene, Antwerp, Belgium. The colony was kept at 25-26°C, 75% relative humidity and 12 hours light/dark photoperiod. A 30% sugar source was permanently provided to adult sand flies. *Phlebotomus pedifer* sand flies were captured in a previous study in Ochollo (6° 11’ N, 37° 41’ E), a village in southwestern Ethiopia where CL is endemic [24,25]. Sand flies were captured between March 2017 and February 2018 using CDC light traps and sticky traps. Specimens were stored in 97% ethanol at −20°C until nucleic acid isolation was carried out in March 2018. *Leishmania* DNA positive sand flies were all *P. pedifer* infected with *L. aethiopica* [24]. Nucleic acid extracts were stored at −20°C until analysis for the current study.

### Hyraxes

Hyraxes had been captured in Ochollo using traditional trapping methods in 2017. Nose and ear samples were collected and stored in 97% ethanol at −20°C until further handling. Molecular analyses revealed that all hyraxes were *H. brucei* infected with *L. aethiopica*. The original tissue samples in 97% ethanol were stored at −20°C until the current analysis [24].

### Assay comparison on field and laboratory infected sand flies and hyraxes

#### Experimental sand fly infection

Laboratory reared *L. longipalpis* were starved 12 hours prior to experimental infection. About 150 sand flies were infected through a chick-skin membrane on heparinized (100 U/mL blood) heat-inactivated mouse blood spiked with *L. major* procyclic promastigotes (5 × 10^6^ promastigotes/mL blood). Engorged females were separated 24 hours post blood meal and were continuously provided with 30% sugar solution. Sand flies were collected six days after infection for dissection of thorax and abdomen (n = 96).

#### Sand fly nucleic acid isolation and purification

Nucleic acids of experimentally infected (*L. major*) *L. longipalpis* were isolated with a crude extraction buffer and purified using an ethanol precipitation approach as described previously [24]. In short, individual sand fly specimens were incubated overnight in 50 µL extraction buffer (10 mM Tris-HCl pH 8, 10 mM EDTA, 0.1 % SDS, 150 mM NaCl) and 0.5 µL proteinase K (200 µg/mL) without maceration. The next day, 25 µL nuclease free water was added and the samples were heated for 5 minutes at 95°C. For nucleic acid precipitation, 20 µL of the extract was supplemented with 1/10^th^ volume 3 M NaOAc (pH 5.6) and 2 volumes 97% ethanol (chilled at −20°C). This suspension was left overnight, after which the samples were centrifuged for 15 minutes at 21,000×*g* at 4°C. The supernatant was removed and 500 µL chilled 70% ethanol was added, followed by centrifugation under the same conditions. The supernatant was removed and the pellet was air-dried for 15 minutes in a heating block at 50°C followed by resuspension in 20 µL nuclease free water.

Additionally, 37 *P. pedifer* nucleic acid extracts were selected from our previous study, of which 17 were identified as *L. aethiopica* positive (kDNA and ITS-1) and 20 as negative (kDNA).

#### DNA/RNA extraction from hyrax samples

Seven *L. aethiopica* DNA positive (kDNA and ITS-1) and 15 negative hyrax tissue samples were selected from our previous study [24]. DNA and RNA were simultaneously extracted from the original tissue samples with the NucleoSpin RNA kit and additional reagents from the NucleoSpin RNA/DNA buffer set (Macherey Nagel, Germany) according to the manufacturer’s instructions.

#### Molecular screening

Nucleic acid extracts of the sand fly and hyrax samples were subjected to three different real-time PCR approaches targeting: *(i)* kDNA/18S DNA in a multiplex Taqman probe assay (further referred to as ‘MP kDNA qPCR’ and ‘MP 18S qPCR’ respectively), *(ii)* kDNA in a SYBR Green assay with an alternate set of primers (‘JW kDNA qPCR’), and *(iii)* SL-RNA in a SYBR Green assay (‘SL-RNA qPCR’). The primers for the JW kDNA qPCR were adopted from Nicolas *et al.* and the assay was carried out as explained in our previous study [14,24], while the other assays were performed as described by Eberhardt *et al.* [23]. All extracts were 1:10 diluted to prevent qPCR inhibition and were run on a Step One Plus real-time qPCR system (Applied Biosystems, Life Technologies, Belgium). The threshold was set at 1 for each qPCR.

### Copy number and extraction method comparison

#### Promastigote isolation from sand fly midgut and culture

We assessed whether there is a copy number difference of kDNA and SL-RNA between parasites isolated from sand fly midguts and *in vitro* cultures in HOMEM. First, an experimental infection of *L. longipalpis* with *L. major* was done to harvest promastigotes from sand fly midguts. About 200 sand flies were collected six days after feeding on a parasitized blood meal and the midguts were dissected under a dissection microscope. Pools of midguts were macerated with a pestle in 100 µL DPBS (Gibco, ThermoFisher Scientific, Belgium) to release the parasites. Second, *L. major* promastigotes from a culture were counted using a KOVA chamber to determine the parasite concentration. An excess volume was taken for further washing steps. Both suspensions were washed twice in 100 µL DPBS with intermediate centrifugation steps of one minute at 21,300×*g*. The pellet was dissolved in 100 µL DPBS. Parasite concentrations were determined in a KOVA chamber and used to prepare two replicates of 10^6^, 10^5^ and 10^4^ parasites in 20 µL DPBS from promastigotes isolated from the sand fly midguts and from culture.

#### DNA/RNA isolation and molecular screening

To determine whether the crude extraction buffer in combination with ethanol precipitation is suitable for efficient nucleic acid isolation and subsequent downstream RNA and DNA detection, nucleic acids from three different concentrations of promastigotes, isolated from either sand fly midguts or culture medium, were extracted using *(i)* a Nucleospin RNA kit and additional RNA/DNA buffer set and *(ii)* the crude extraction buffer and ethanol precipitation approach. For the latter, the complete volume of the nucleic acid extract was used for ethanol precipitation. The final elution volumes were equalized to ensure the same relative DNA and RNA yields for both methods. All extracts were subjected in duplicate to the JW kDNA and SL-RNA qPCRs.

### Sand fly spiking

Promastigotes (*L. major*) were harvested from a stationary-phase culture and washed with DPBS. The number of promastigotes was determined in a KOVA counting chamber and the pellet was stored at - 20°C until extraction. Naive *L. longipalpis* sand flies were spiked with a 10-fold serial dilution of *L. major* promastigotes, ranging from 1.6 × 10^7^ to 1.6 × 10^−6^ parasites. The samples were extracted with the crude extraction buffer and ethanol precipitation approach, and subsequently subjected in duplicate to the SL-RNA qPCR.

### Data analysis

Analyses were carried out using GraphPad Prism version 8 (GraphPad Software, La Jolla California, USA). The correlation between the Ct values of the SL-RNA assay and the other three qPCRs was determined by a Pearson correlation. This analysis was performed using the infected field-collected sand flies because of the broad range of Ct values. A standard curve with linear regression and PCR efficiency was generated to determine the theoretical LoD and limit of quantification (LoQ) of the SL-RNA qPCR.

## Results

### Comparison of *Leishmania* detection assays

Of the 96 *L. major* infected laboratory *L. longipalpis* sand flies, two samples were negative and 82 were positive by all assays (Fig 1A, S1 Table A). Among the samples that were positive by all qPCRs, the 18S DNA qPCR showed the highest mean Ct value (30.3 ± 2.3), followed by the MP kDNA qPCR (17.3 ± 1.4), JW kDNA qPCR (14.6 ± 1.4) and SL-RNA qPCR (13.8 ± 0.9). Ten samples were not detected with the 18S DNA qPCR, but were positive for the other three assays. These samples had higher mean Ct values of 21.4 (± 4.4), 17.5 (± 3.1) and 17.1 (± 2.4) for the MP kDNA, JW kDNA and SL-RNA assays respectively. Two sand fly specimens with the highest Ct values in the JW kDNA (27.1 ± 0.2) and SL-RNA qPCRs (25.7 ± 0.7) could not be identified by the MP kDNA qPCR. Overall, the JW kDNA and SL-RNA qPCRs provided concordant results and performed equally well on the laboratory infected sand flies.

**Fig 1:**
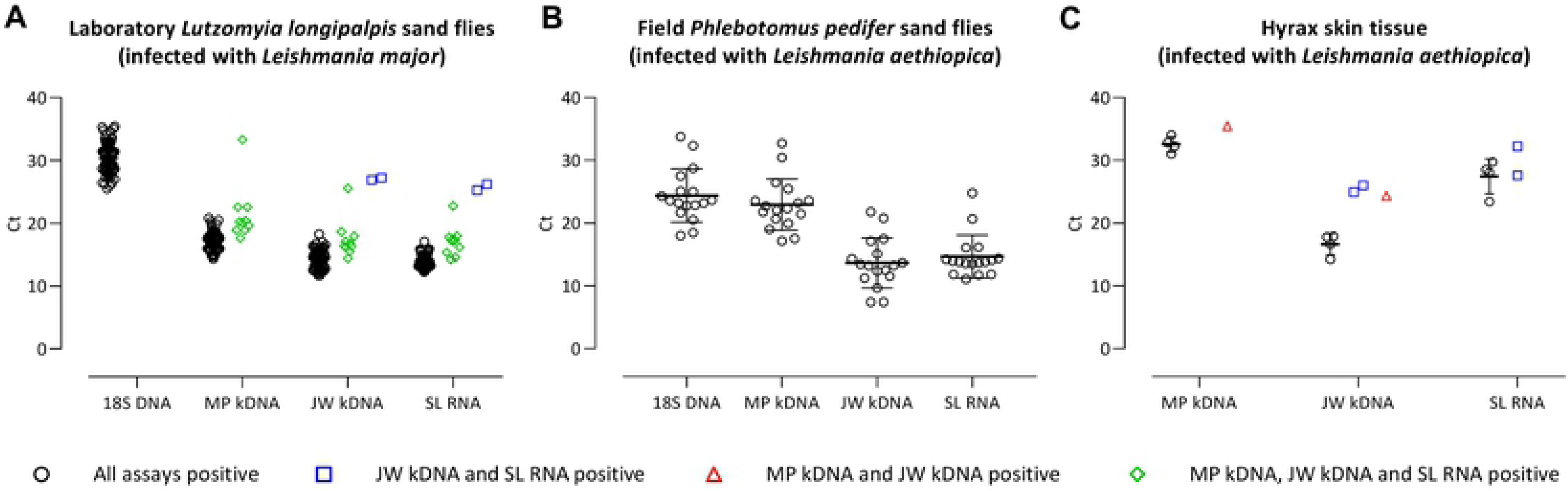
PCR cycle threshold (Ct) values of laboratory and field infected sand flies and infected hyrax tissue samples. Mean Ct value and error bars (standard deviation) are presented for the samples that were positive in all assays that they were tested for. Due to technical issues, the analysis of the 18S DNA qPCR on hyrax tissue samples was not included.

Among the field collected, ethanol stored sand fly specimens, 20 were negative and 17 positive in all four assays (Fig 1B, S1 Table B). Mean Ct values of the JW kDNA and SL-RNA qPCRs were similar (13.7 ± 3.9 and 14.7 ± 3.4 respectively) and consistently lower than the Ct values obtained by the other two assays (18S DNA: 24.2 ± 4.2 and MP kDNA: 22.9 ± 4.2). The difference in Ct values between the MP kDNA and JW kDNA qPCRs was larger for *P. pedifer* infected *with L. aethiopica* than for *L. longipalpis* infected with *L. major* (Fig 1A versus 1B).

Seven out of 22 long-term stored hyrax tissue samples tested positive in two or more qPCR assays (Fig 1C, S1 Table C). Four samples were positive in all assays, resulting in the lowest Ct values for the JW kDNA qPCR (16.6 ± 1.7), compared to the SL-RNA qPCR (27.5 ± 2.8) and MP kDNA qPCR (32.6 ± 1.2). Two samples with high Ct values in the JW kDNA and SL-RNA qPCRs were negative for the MP kDNA qPCR, while one sample was positive for the MP kDNA and JW kDNA qPCRs with high Ct values, but not for the SL-RNA assay.

Overall, the Pearson correlation showed that the SL-RNA qPCR correlated quite well with the Ct values of the JW kDNA (Fig 2A; R^2^ = 0.82, n = 17), MP kDNA (Fig 2B; R^2^ = 0.90, n = 17) and 18S DNA assays (Fig 2C; R^2^ = 0.88, n = 17). For all comparisons, the confidence intervals increased towards the higher Ct values, which could be due to slight inhibition of the SL-RNA qPCR.

**Fig 2:**
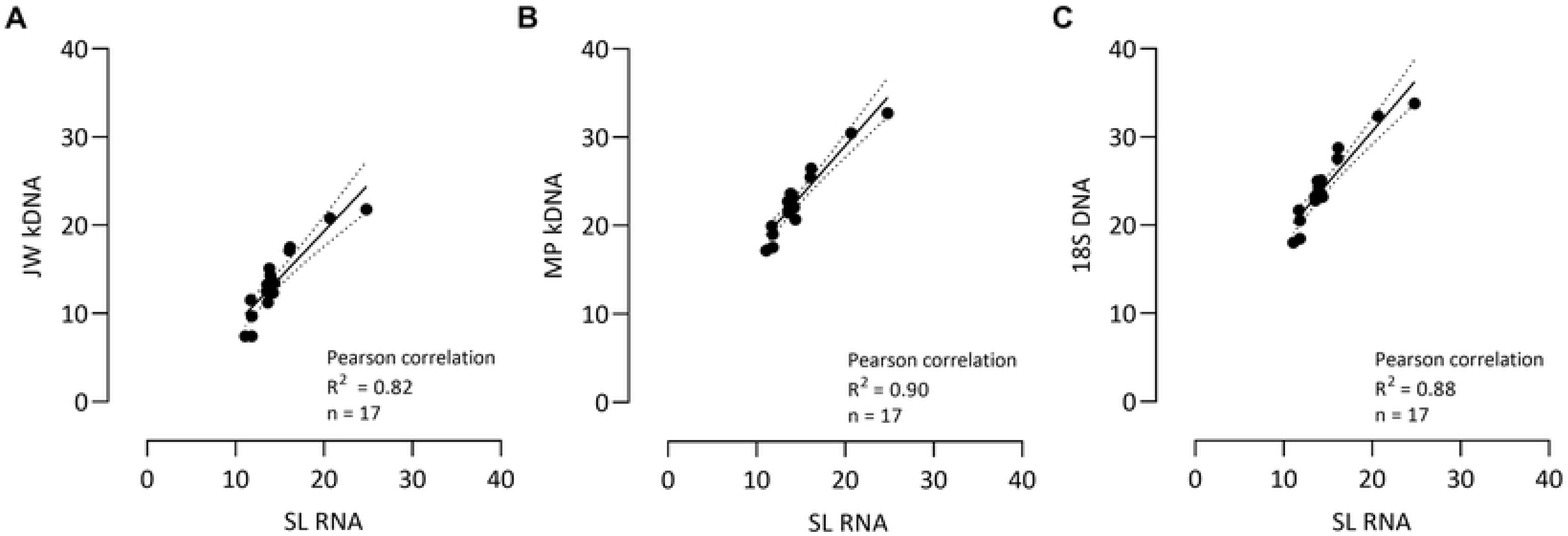
Correlation between Ct values obtained by the SL-RNA qPCR and the (A) JW kDNA qPCR, (B) MP kDNA qPCR and (C) 18S DNA qPCR. Pearson correlation analysis of the results obtained with the different assays. Linear regression and 95% confidence intervals (dotted lines) are shown in the graphs. Ct = cycle threshold value.

### Extraction method comparison and copy number difference

The crude extraction buffer with ethanol precipitation and column purification (respectively referred to as ‘crude’ and ‘column’ in Fig 3) methods showed similar extraction efficiencies for kDNA, with comparable Ct values obtained for the standardized concentrations of promastigotes isolated from culture or sand fly midguts. Likewise, both methods performed well for SL-RNA extraction, although the RNA yield appeared even slightly higher (on average 1.5 Ct lower values) with the crude method. The kDNA copy number was similar for promastigotes isolated from culture and sand fly midguts (Fig 3, grey versus black symbols). For SL-RNA, both extraction methods revealed that the Ct values for sand fly derived promastigotes were slightly but consistently higher (1.7 Ct on average) than those for culture-derived promastigotes. The JW kDNA qPCR reaction suffered inhibition in both runs for 10^6^ promastigotes isolated from sand fly midguts (Fig 3, lacking grey circle for ‘crude’).

**Fig 3:**
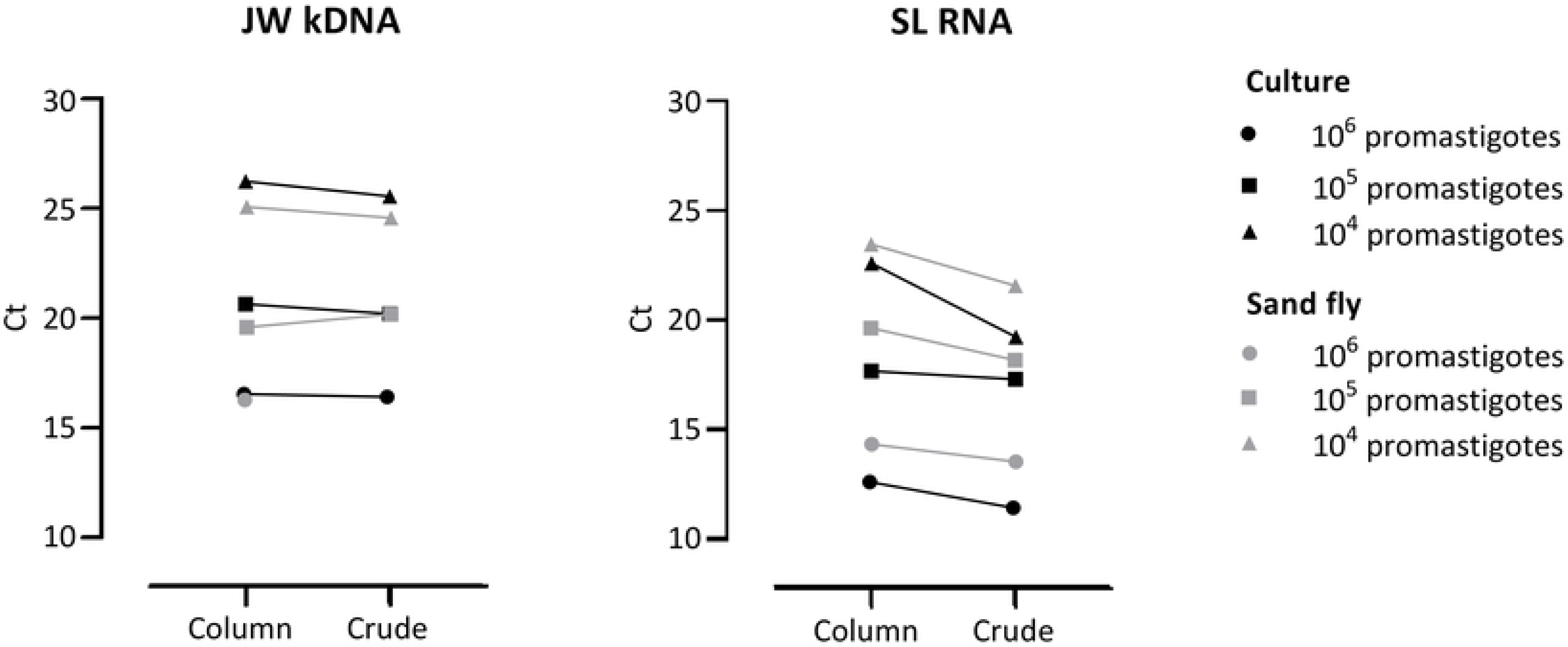
Extraction method and copy number comparison. Ct values of promastigotes isolated from culture (black symbols) and sand fly midguts (grey symbols) that were extracted with a commercial column extraction (‘column’) or crude high-salt extraction buffer (‘high-salt’) and subjected to JW kDNA and SL-RNA qPCRs. Each symbol presents the assay result for a standardized concentration of promastigotes that was used for the comparisons. Ct = cycle threshold value.

### LoD and LoQ of the SL-RNA qPCR

Based on the serial dilution of *L. longipalpis* sand flies spiked with *L. major* promastigotes, the theoretical LoD of the SL-RNA qPCR was 10^−3^ parasite equivalents (Fig 4A). For 1.6 × 10^7^ promastigotes, the assay did not provide a result in any of the two independent runs, implying that there was PCR inhibition at this concentration. The assay showed a very good PCR efficiency of 105% for the serial dilution down to 10 parasites, representing the theoretical LoQ. A Pearson correlation demonstrated an excellent inter-run stability for the two independent runs of the SL-RNA qPCR on the serial dilution (Fig 4B; R^2^ = 0.99, n = 10).

**Fig 4:**
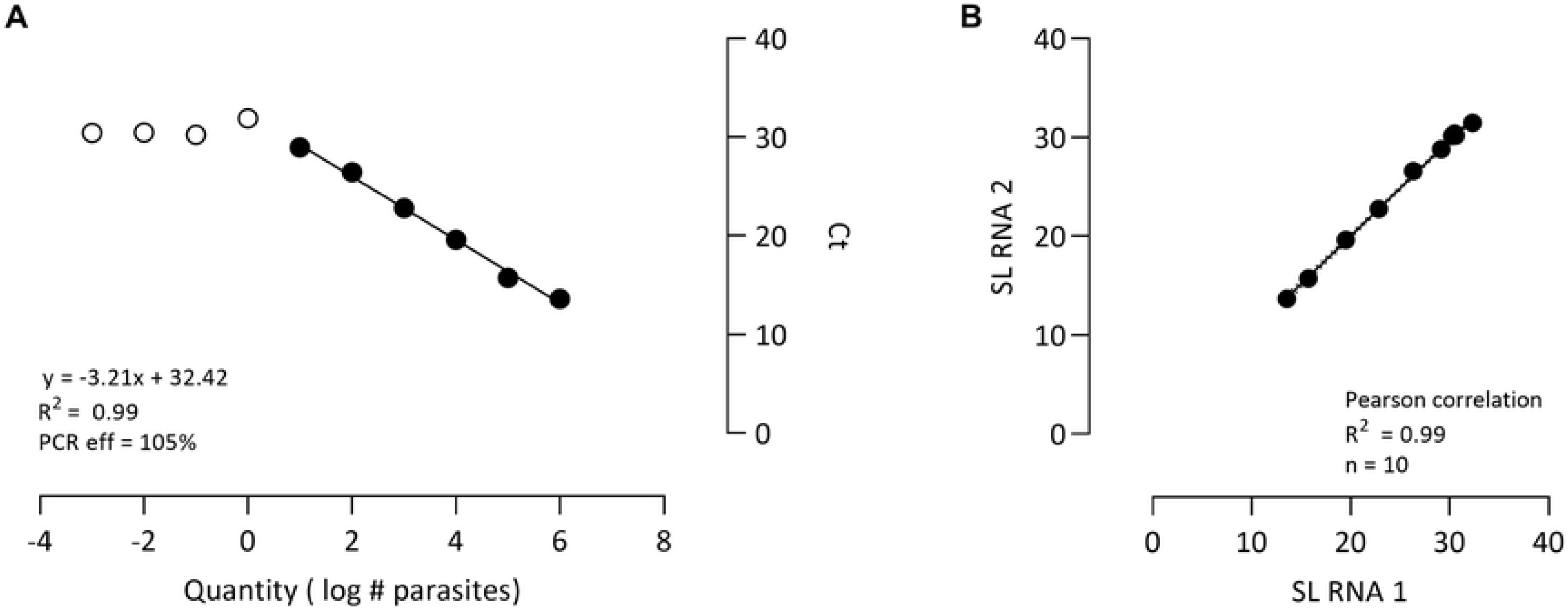
Performance of the SL-RNA qPCR on a serial dilution of *L. longipalpis* sand flies spiked with *L. major* promastigotes. (A) Standard curve with linear regression and qPCR efficiency. The open symbols depict all concentrations that were detected by the assay, while the filled symbols are the parasite concentrations that show a linear correlation. (B) Inter-run variability of the SL-RNA qPCR in two replicates, analyzed by a Pearson correlation and linear regression analysis. Dotted lines indicate the 95% confidence intervals. Ct = cycle threshold value.

## Discussion

For eco-epidemiological surveys, researchers currently opt for sensitive high-throughput molecular screening methods for *Leishmania* detection in sand flies and potential reservoirs, because infection prevalence is overall quite low, even in endemic areas [24,26]. These molecular methods most often target DNA sequences, which may persist for quite some time after parasite death. Hosts and sand flies can be exposed to the parasite without establishment of an infection. Hence, it would be highly informative to be able to specifically detect viable parasites. In our study, we evaluated whether the recently developed SL-RNA qPCR assay by Eberhardt *et al.* [23] enables *Leishmania* detection in sand flies and skin tissue from CL infected animals. The targeted 39 bp SL-RNA sequence is conserved amongst *Leishmania* species and fulfils an essential function in RNA trans-splicing and polyadenylation processes [27]. To our knowledge, this study is the first to evaluate the use of an RNA target for *Leishmania* detection in vectors and field-sampled tissue of reservoir hosts. Since RNA quickly decays after death of the infectious agent, it is considered as a promising detection marker for viable *Leishmania* parasites [22,28,29], although the half-life of SL-RNA as small nuclear RNA molecule remains to be determined. We comparatively assessed the performance of four molecular diagnostic assays on field and laboratory infected sand flies and hyrax tissue samples. The JW kDNA [14,24] and SL-RNA qPCRs [23] showed the best performance using the laboratory *L. major*-infected *L. longipalpis* and field-collected *L. aethiopica*-infected *P. pedifer* sand flies. Both assays identified the same positive and negative samples, indicating that they have a similar analytical sensitivity and specificity. It was surprising that the SL-RNA qPCR performed very well on field-collected sand flies, considering that these samples had not been preserved under favorable conditions for RNA, which may relate to the short amplicon length [30,31]. These observations indicate that SL-RNA could be an interesting target for *Leishmania* detection in vectors collected during entomological surveys.

On the contrary, the MP kDNA and 18S DNA qPCRs could not identify all positive laboratory infected sand flies. One reason is the low copy number of the 18S rDNA fragment (50-200 copies per *Leishmania* genome) [32] compared to the much higher copy number of SL-RNA (a single *Trypanosoma* cell contains about 8,600 copies [33]) and kDNA (a *Leishmania* parasite contains approximately 10,000 copies [34]). Whereas the copy number of kDNA was fairly similar in promastigotes isolated from sand flies and from culture, SL-RNA appears to be slightly less abundant in parasites isolated from sand flies. This may indicate a reduced transcriptional activity of the vector-derived parasite pool (containing various life cycle stages) as compared to culture-derived parasites.

The MP kDNA qPCR could not identify some of the laboratory infected sand flies that were positive by the JW kDNA and SL-RNA qPCRs, most probably because multiplex qPCRs are commonly slightly less sensitive than uniplex assays [35,36]. Additionally, the MP kDNA qPCR provided higher Ct values on the field-collected sand flies, which most likely relates to mismatches of the reverse primer with the *L. aethiopica* kDNA fragment (S1 Fig) [17]. Earlier observations of a lower sensitivity for *L. tropica* (genetically very similar to *L. aethiopica*) and *L. mexicana* [23] corroborates the limitations of this MP kDNA assay that was originally developed by Mary *et al.* for detection of *L. donovani* [17]. The SL-RNA qPCR provided equal Ct values for various *Leishmania* species, demonstrating its suitability as a pan-*Leishmania* assay [23].

Although only a few positive hyrax tissue samples were tested, the JW kDNA qPCR performed best under the used sample storage conditions. A sample was considered positive if identified by two different assays. The SL-RNA assay identified one false negative sample and showed generally higher Ct values compared to the JW kDNA PCR than for the sand fly screening, which is probably due to the fact that the samples had been stored in ethanol for two years before DNA/RNA extraction was performed. Most likely, proper RNA storage conditions and/or immediate RNA isolation would result in a substantially improved performance of the SL-RNA qPCR on tissue samples [31,33]. Favoring this viewpoint, the SL-RNA qPCR showed excellent analytical sensitivity in laboratory infected (*L. infantum*) mouse spleen and liver samples, detecting down to 10^−3^ parasite equivalents per mg tissue [23].

Considering the large sample size that needs to be screened in search for positive field specimens, a low-cost, efficient nucleic acid extraction method is preferred [9]. We found that a crude extraction buffer in combination with an ethanol precipitation step is as efficient as a commercial column extraction for downstream DNA and RNA detection in sand flies. Ct values tended even slightly lower when the extraction was carried out with the crude method, suggesting that there is some nucleic acid loss on the silica columns. Other important advantages of this crude extraction method are the low-cost and reduction in sample processing time as maceration is not required [8,9]. The latter is compensated by a more time-consuming ethanol precipitation step. However, because of the low prevalence in field collected sand flies, individual extracts can be pooled to reduce the number of samples for purification and PCR [24].

Determination of the viable parasite load in sand flies can be highly informative, especially for studies that investigate *e.g.* the vectorial capacity. Previously, the LoD of the SL-RNA qPCR on cultured promastigotes has been established at 0.0002 parasite equivalents [23]. We assessed the theoretical LoD of the SL-RNA qPCR based on sand flies spiked with a serial dilution of *L. major* promastigotes. The determined theoretical LoD of 10^−3^ parasite equivalents per reaction of our assay is similar to findings of Bezerra-Vasconcelos *et al.*, who could detect 10^−3^ parasites per reaction with a kDNA qPCR assay on *L. infantum*-spiked *L. longipalpis* sand flies [13]. This substantiates that the sensitivity of qPCR assays targeting SL-RNA and kDNA are comparable, which corroborates the comparative assessment performed in this study. Moreover, congruence of the assays appears very good, indicating that both can achieve reliable quantification. Based on the standard curve, it can be concluded that SL-RNA qPCR can quantify down to 10 parasites per sand fly with high PCR efficiency, which is sufficient for determination of biologically relevant parasite loads.

Overall, this study shows for the first time that the SL-RNA target can be used for detection and quantification of *Leishmania* parasites in field and laboratory infected sand flies, even in combination with a crude extraction method. The SL-RNA qPCR assay provides complementary information to the standard kDNA assays, as it is pan-*Leishmania* specific and able to detect viable parasites, which can be a major advantage for eco-epidemiological studies including identification of vectors and reservoirs.

## Acknowledgements

We are grateful to Dr. Gert Van der Auwera (Institute of Tropical Medicine, Antwerp, Belgium) for his advice on the interpretation of the experiments. We would also like to thank the village head and field workers in Ochollo for the possibility to collect the sand fly and hyrax samples.

## Supporting information captions

**S1 Table: qPCR results of sand fly and hyrax samples from the laboratory and field in four different assays.** For each assay the mean cycle threshold value (± standard deviation) are presented.

**S1 Fig: Annealing of MP kDNA qPCR primers to *L. aethiopica* kDNA (GenBank: U77892.1).**

**S1 Data: Dataset of laboratory infected (*Leishmania major*) *Lutzomyia longipalpis* sand flies screened by SL-RNA, 18S DNA, MP kDNA and JW kDNA qPCR assays.**

**S2 Data: Dataset of field collected (Ethiopia) *Phlebotomus pedifer* sand flies (some infected by *Leishmania aethiopica*) screened by SL-RNA, 18S DNA, MP kDNA and JW kDNA qPCR assays.**

**S3 Data: Dataset of Dataset of field (Ethiopia) collected *Heterohyrax brucei* tissue samples (some infected by *Leishmania aethiopica*) screened by SL-RNA, 18S DNA, MP kDNA and JW kDNA qPCR assays.**

**S4 Data: Data of extraction method and copy number comparison.**

**S5 Data: Data of standard curve of the SL-RNA qPCR based on *Lutzomyia longipalpis* sand flies spiked with *Leishmania major*.**

## References

1. Parasites - Leishmaniasis [Internet]. Center of disease control and prevention. 2013 [cited 2018 Sep 7]. Available from: https://www.cdc.gov/parasites/leishmaniasis/biology.html

2. Bray RS, Ashford RW, Bray MA. The parasite causing cutaneous leishmaniasis in Ethiopia. Trans R Soc Trop Med Hyg. 1973;67(3):345–8.

3. Ashford W, Bray M, Hutchinson P, Bray S. The epidemiology of cutaneous leishmaniasis in Ethiopia. Trans R Soc Trop Med Hyg. 1973;67(4).

4. Lemma W, Erenso G, Gadisa E, Balkew M, Gebre-michael T, Hailu A. A zoonotic focus of cutaneous leishmaniasis in Addis Ababa, Ethiopia. Parasit Vectors. 2009;2(60).

5. Ashford RW. A possible reservoir for *Leishmania tropica* in Ethiopia. Trans R Soc Trop Med Hyg. 1970;64(6):936–7.

6. Rogers ME, Bates PA. *Leishmania* Manipulation of Sand Fly Feeding Behavior Results in Enhanced Transmission. PLOS Pathog. 2007;3(6):1–7.

7. Galluzzi L, Ceccarelli M, Diotallevi A, Menotta M, Magnani M. Real-time PCR applications for diagnosis of leishmaniasis. Parasit Vectors. 2018;11(273).

8. Asghar U, Malik MF, Anwar F, Javed A, Raza A. DNA Extraction from Insects by Using Different Techniques: A Review. Adv Entomol. 2015;3:132–8.

9. Kato H, Uezato H, Gomez EA, Terayama Y, Calvopiña M, Iwata H. Establishment of a mass screening method of sand fly vectors for *Leishmania* infection by molecular biological methods. Am J Trop Med Hyg. 2007;77(2):324–9.

10. Odiwuor SOC, Saad AA, Doncker S De, Maes I, Laurent T. Universal PCR assays for the differential detection of all Old World Leishmania species. 2011;209–18.

11. Simpson L, Douglass SM, Lake JA, Pellegrini M, Li F. Comparison of the Mitochondrial Genomes and Steady State Transcriptomes of Two Strains of the Trypanosomatid Parasite, Leishmania tarentolae. PLoS Negl Trop Dis. 2015;9(7):1–35.

12. Aransay ANAM, Scoulica E, Tselentis Y. Detection and Identification of Leishmania DNA within Naturally Infected Sand Flies by Seminested PCR on Minicircle Kinetoplastic DNA. 2000;66(5):1933–8.

13. Bezerra-vasconcelos DR, Melo LM, Albuquerque ÉS, Luciano MCS, Bevilaqua CML. Real-time PCR to assess the *Leishmania* load in *Lutzomyia longipalpis* sand flies: Screening of target genes and assessment of quantitative methods. Exp Parasitol. 2011;129:234–9.

14. Nicolas L, Prina E, Lang T, Milon G. Real-Time PCR for Detection and Quantitation of Leishmania in Mouse Tissues. 2002;40(5):1666–9.

15. Kassahun A, Sadlova J, Dvorak V, Kostalova T, Rohousova I, Frynta D, et al. Detection of *Leishmania donovani* and *L. tropica* in Ethiopian wild rodents. Acta Trop. 2015;145:39–44.

16. Kassahun A, Sadlova J, Benda P, Kostalova T, Warburg A, Hailu A, et al. Natural infection of bats with *Leishmania* in Ethiopia. Acta Trop. 2015;150:166–70.

17. Mary C, Lascombe L, Dumon H, Al MET, Icrobiol JCLINM. Quantification of *Leishmania infantum* DNA by a Real-Time PCR Assay with High Sensitivity. J Clin Microbiol. 2004;42(11):5249–55.

18. Weirather JL, Jeronimo SMB, Gautam S, Sundar S, Kang M, Kurtz MA, et al. Serial Quantitative PCR Assay for Detection, Species Discrimination, and Quantification of *Leishmania spp.* in Human Samples. J Clin Microbiol. 2011;49(11):3892–904.

19. Volpini C, Passos VMA, Correa G, Romanha AJ. PCR-RFLP to identify Leishmania (Viannia) braziliensis and L. (Leishmania) amazonensis causing American cutaneous leishmaniasis. 2004;90:31–7.

20. Boni SM, Oyafuso LK, Soler RDC. Efficiency of noninvansive sampling methods (swab) together with Polymerase Chain Reaction (PCR) for diagnosing American Tegumentary Leishmaniasis. Rev do Inst Med Trop Sao Paulo. 2017;

21. Satow MM, Yamashiro-kanashiro EH, Rocha MC, Oyafuso LK, Soler RC. Applicability of kDNA-PCR for routine diagnosis of American tegumentary leishmaniasis in a tertiary reference hospital. 2013;55(6):393–9.

22. Keer JT, Birch L. Molecular methods for the assessment of bacterial viability. J Microbiol Methods. 2003;53:175–83.

23. Eberhardt E, Van Den Kerkhof M, Bulté D, Mabille D, Van Bockstal L, Monnerat S, et al. Evaluation of a pan-*Leishmania* Spliced-Leader RNA detection method in human blood and experimentally infected syrian golden hamsters. J Mol Diagnostics. 2018;20(2).

24. Pareyn M, Van Den Bosch E, Girma N, Houtte N Van, Van Dongen S, Van Der Auwera G, et al. Ecology and seasonality of sandflies and potential reservoirs of cutaneous leishmaniasis in Ochollo, a hotspot in southern Ethiopia. PLoS Negl Trop Dis. 2019;13(8).

25. Mengistu G, Laskay T, Gemetchu T, Humber D, Ersamo M, Eva D, et al. Cutaneous leishmaniasis in south-western Ethiopia: Ocholo revisited. Trans R Soc Trop Med Hyg. 1992;86:149–53.

26. Assefa A. Leishmaniasis in Ethiopia : A systematic review and meta-analysis of prevalence in animals and humans. Heliyon. 2018;(June):e00723.

27. Perry KL, Watkins KP, Agabian N. Trypanosome mRNAs have unusual “cap 4” structures acquired by addition of a spliced leader. Proc Natl Acad Sci USA. 1987;84:8190–4.

28. Analysis RP, Cangelosi GA, Weigel KM, Lefthand-begay C, Meschke JS. Molecular Detection of Viable Bacterial Pathogens in Water by Ratiometric Pre-rRNA Analysis. Appl Environ Microbiol. 2010;76(3):960–2.

29. Gabaint AVON, Belasco JG, Schottel JL, Annie CY, Cohen SN. Decay of mRNA in *Escherichia coli*: Investigation of the fate of specific segments of transcripts. Natl Acad Sci. 1983;80:653–7.

30. Debode F, Marien A, Janssen É, Bragard C, Berben G. The influence of amplicon length on real-time PCR results. Biotechnol Agron Soc Environ. 2017;21(1):3–11.

31. Kuang J, Yan X, Genders AJ, Granata C, Bishop DJ. An overview of technical considerations when using quantitative real-time PCR analysis of gene expression in human exercise research. PLoS One. 2018;13(5).

32. Srivastava P, Mehrotra S, Tiwary P, Chakravarty J, Sundar S. Diagnosis of Indian Visceral Leishmaniasis by Nucleic Acid Detection Using PCR. PLoS One. 2011;6(4).

33. González-andrade P, Camara M, Ilboudo H, Bucheton B, Jamonneau V. Diagnosis of Trypanosomatid Infections Targeting the Spliced Leader RNA. J Mol Diagnostics. 2014;16(4):400–4.

34. Simpson L. Kinetoplast DNA in Trypanosomid Flagellates. Int Rev Cytol. 1986;99:119–79.

35. Tuo D, Shen W, Yang Y, Yan P, Li X, Zhou P. Development and Validation of a Multiplex Reverse Transcription PCR Assay for Simultaneous Detection of Three Papaya Viruses. Viruses. 2014;341:3893–906.

36. Yao B, Wang G, Ma X, Liu W, Tang H, Zhu H. Simultaneous detection and differentiation of three viruses in pear plants by a multiplex RT-PCR. J Virol Methods. 2014;196:113–9.

